# New antiviral defences are genetically embedded within prokaryotic immune systems

**DOI:** 10.1101/2024.01.29.577857

**Authors:** Leighton J. Payne, Tom C. D. Hughes, Peter C. Fineran, Simon A. Jackson

## Abstract

Bacteria and archaea typically have multiple defence systems that protect them against viral predation. Recently, many new defence systems have been discovered, yet the full scope of the prokaryotic pan-immune system remains to be determined. In this study, we observed that many multi-gene defence systems have additional genes nested or ‘embedded’ within them. Based on this observation, we present a new approach to predict new defence systems, where defence function of uncharacterised genes is inferred based on their genetic embedding in known defence systems. Applying this ‘guilt-by-embedding’ method, we identified and confirmed anti-phage function for seven defence systems and predicted 145 additional candidates. Our findings expand the known immune repertoire of prokaryotes, provide a wealth of new systems for future functional studies, and demonstrate a simple, efficient approach to identify new antiviral defences.

## INTRODUCTION

Diverse antiviral defence systems have evolved in bacteria and archaea that defend them against phages, archaeal viruses and other mobile genetic elements (MGEs) (Georjon and Bernheim, 2023; Mayo-Muñoz et al., 2023). The protein domains associated with defence are wide-ranging and are often also found in other conserved cellular systems, complicating *in silico* discovery of new defence systems based on sequence alone. To address this issue, contextual genetic analyses have been used in defence system discovery with great success. Applying the principle of ‘guilt-by-association’ (Aravind, 2000a; Galperin and Koonin, 2000; Huynen et al., 2000) to identify genes functionally linked to prokaryotic Argonautes led to the discovery of defence islands—genomic regions of colocalised defence systems (Makarova et al., 2009, 2011, 2013). The defence island concept inspired the systematic prediction of new systems by investigating protein domains or families enriched in the proximity of known defence systems, greatly expanding our view of the prokaryotic pan-immune system (Goldfarb et al., 2015; Ofir et al., 2017; Doron et al., 2018; Bernheim and Sorek, 2020; Gao et al., 2020; Bernheim et al., 2021; Garb et al., 2022; LeRoux et al., 2022; Millman et al., 2022). However, an alternative high-throughput functional screening approach to system discovery demonstrated that there are likely many uncharacterised defence systems that have not been predicted by defence island enrichment methods (Vassallo et al., 2022).

Here, we introduce a new approach to system discovery, which we have termed ‘guilt-by-embedding’, where defence function is inferred for uncharacterised genes based on their genetic embedding in known defence systems. By analysing the genes embedded in several DNA modification-based defence systems, we identified and confirmed phage defence function for seven systems, and predicted 145 additional candidates. We show that many of these new systems are not typically genetically enriched near other defence systems, demonstrating that our approach can identify systems not detected by methods that rely on defence island enrichment scores. Overall, our findings expand the known defence repertoire of prokaryotes and demonstrate a new, simple approach to identify defence system candidates for functional testing.

## RESULTS

### Antiviral defence systems are frequently embedded in Hma systems

Previously, we discovered Hma (Helicase, Methylase, ATPase) defence systems, which were frequently co-localised with Septu systems (Payne et al., 2021). Closer inspection of this association revealed that Septu was not typically found adjacent to Hma, as would be expected in a typical defence island (Makarova et al., 2011, 2013; Hochhauser et al., 2023). Instead, Septu was commonly encoded between the *hma* genes, an architecture we refer to herein as being ‘embedded’ within Hma (**Figure 1A**). Several other defence systems were also occasionally embedded in Hma systems (**Figure 1A**). To quantify the embedding of defences in Hma, we used PADLOC to search for Hma and other defence systems in more than 230,000 bacterial and archaeal genomes. Of the 1,485 distinct Hma systems identified, 931 (62.7%) contained embedded genes (**Figure 1B**). Hma systems both with or without embedded genes were not typically found near other defence systems (**Figure 1C**). Approximately 22.8% of Hma-embedded genes encoded known defence systems and 3.1% encoded mobility-associated proteins (MAPs) such as transposases, integrases, and phage proteins (**Figure 1D**). Overall, Hma-embedded genes were diverse, encoding 1,555 non-redundant proteins, which we grouped into 434 protein families using sequence clustering and Hidden Markov Model (HMM)-HMM comparisons (**Figure 1E**). Based on these protein family assignments, there were 376 unique Hma-embedded gene cassettes. Since many different known phage defence systems were occasionally embedded in Hma, we hypothesised that many of the Hma-embedded gene cassettes of unknown function would likely include new phage defence systems.

**Figure 1.**
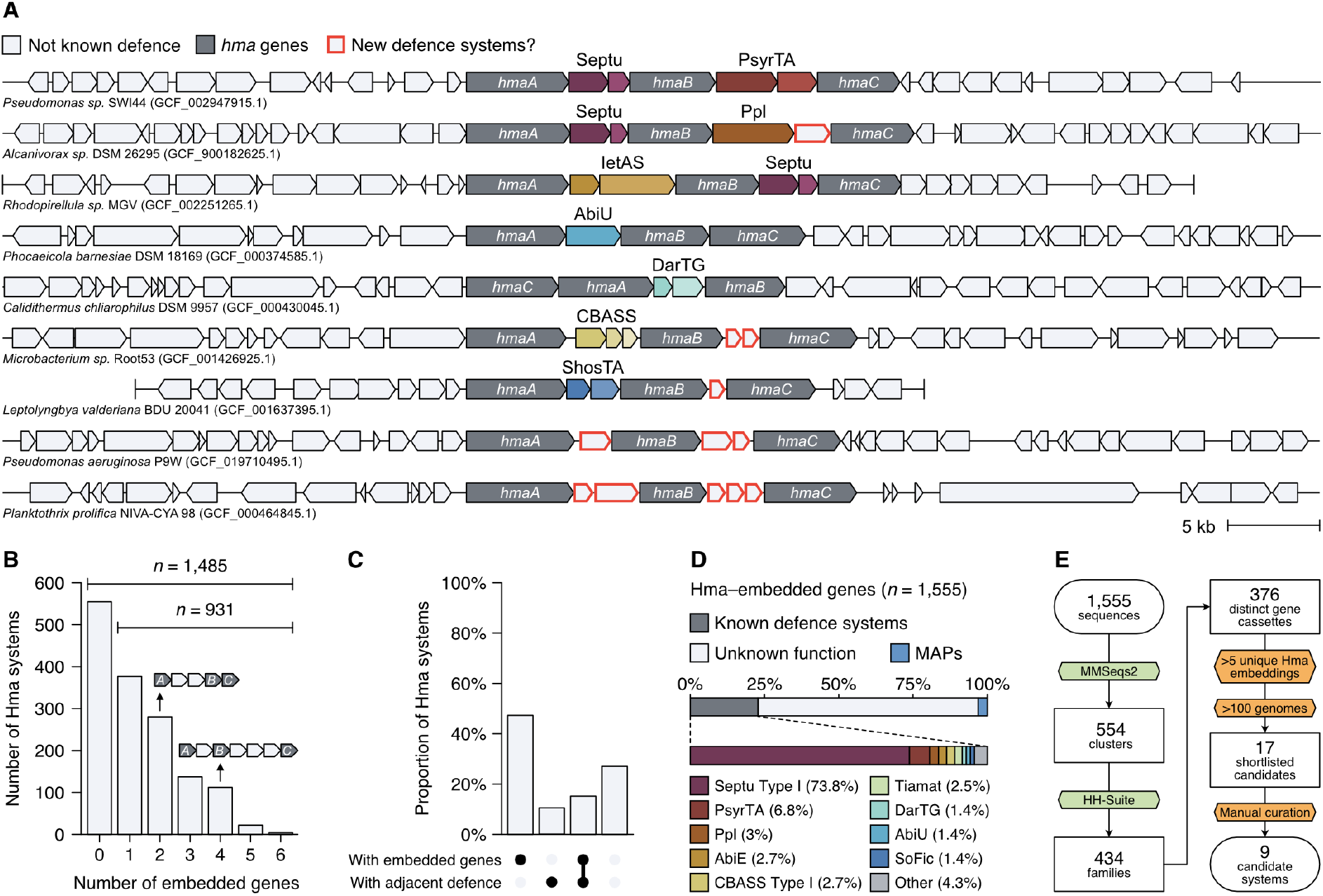
Hma systems often contain additional embedded genes, many of which encode phage defences. (A) Examples of *hma* loci with embedded genes. Vertical bars indicate contig ends. (B) Number of embedded genes identified in Hma systems, allowing for up to three embedded genes between each *hma* gene. Examples of two and four embedded genes are shown. (C) Proportion of Hma systems with and without embedded genes, and with and without adjacent defence systems (up to 10 open reading frames (ORFs) on either side of Hma). (D) Proportion of Hma-embedded genes belonging to known defence systems (grey; further detailed underneath) and mobility-associated proteins (MAPs; blue) or otherwise with unknown function (white). (E) Bioinformatic pipeline for identifying defence system candidates embedded in Hma systems. Tools are highlighted in green and filtering steps are highlighted in orange.

To select a reduced set of defence system candidates for experimental testing, we filtered the uncharacterised gene cassettes for those embedded in more than five Hma systems (**Figure 1E**), then assigned protein domain annotations to each family based on HMM-HMM comparison with Pfam v35 (El-Gebali et al., 2019) (**Table S1**). We expected that phage defence systems capable of functioning independently of Hma would also commonly be found outside of *hma* loci, as is the case for Septu systems, where only 0.7% are embedded in Hma (Payne et al., 2021). Therefore, we built PADLOC models for each defence candidate, searched these models back across the genome dataset, then filtered for systems present in more than 100 genomes, constituting a shortlist of 17 candidate defence systems (**Figure S1**). Most of these candidates were not enriched near other defence systems (**Figure S1**). Based on their apparent novelty, we selected nine candidate systems to test for phage defence. For simplicity herein, the candidate systems are identified with the prefix HEC (<u>H</u>ma-<u>e</u>mbedded <u>c</u>andidate) followed by a unique integer.

### Hma-embedded candidate systems exhibit antiviral activity

The candidate systems encoded diverse protein families, including three ATPases associated with either a topoisomerase-primase (TOPRIM) (HEC-01), nuclease-related (NERD) (HEC-02), or PilT N-terminal (PIN) (HEC-03) domain-containing protein; an ATPase fused to a TOPRIM domain (HEC-04); two GmrSD-like proteins (HEC-05 and HEC-06); and RelE (HEC-07), higher eukaryotes and prokaryotes nucleotide-binding (HEPN) (HEC-08) and Nedd4-BP1/YacP nuclease (NYN) (HEC-09) domain-containing proteins (**Figure 2A; Table S2**). Subsequent to the selection of HEC-05 as a candidate, a highly similar HEC-05 homolog was discovered and characterised as the defence system BrxU (Picton et al., 2021). To assess defence function, one representative of each candidate system (including its putative native promoter region) was cloned into a plasmid that included an upstream inducible promoter to supplement the native promoter activity in *Escherichia coli*. The systems were expressed in *E. coli* K-12 MG1655 ΔRM (Maffei et al., 2021) and challenged with a subset of the BASEL phage collection (Maffei et al., 2021) and several ‘classic’ coliphages. An initial qualitative assessment of plaque reduction, providing resolution to detect at least a 10-fold reduction in efficiency of plaquing (EOP) or a reduction in plaque size compared to a non-defence control (expressing mCherry), revealed seven systems that reduced plaquing of at least one phage (**Figure 2B**). HEC-01 and HEC-09 did not provide protection against any of the phages tested (**Figure 2B**). A quantitative assessment of the remaining systems showed that these were able to reduce the EOP of at least two phages by several orders of magnitude compared to the control (**Figure 2B; Figure S2**). For the two-gene systems HEC-02 and HEC-03, both genes were required for defence (**Figure S3**). Considering that lack of activity against the phages we tested does not rule out a potential function against other phages or different types of MGEs, our validation of seven of the nine systems tested demonstrates the power of our guilt-by-embedding approach to identify phage defence systems with a high success rate.

**Figure 2.**
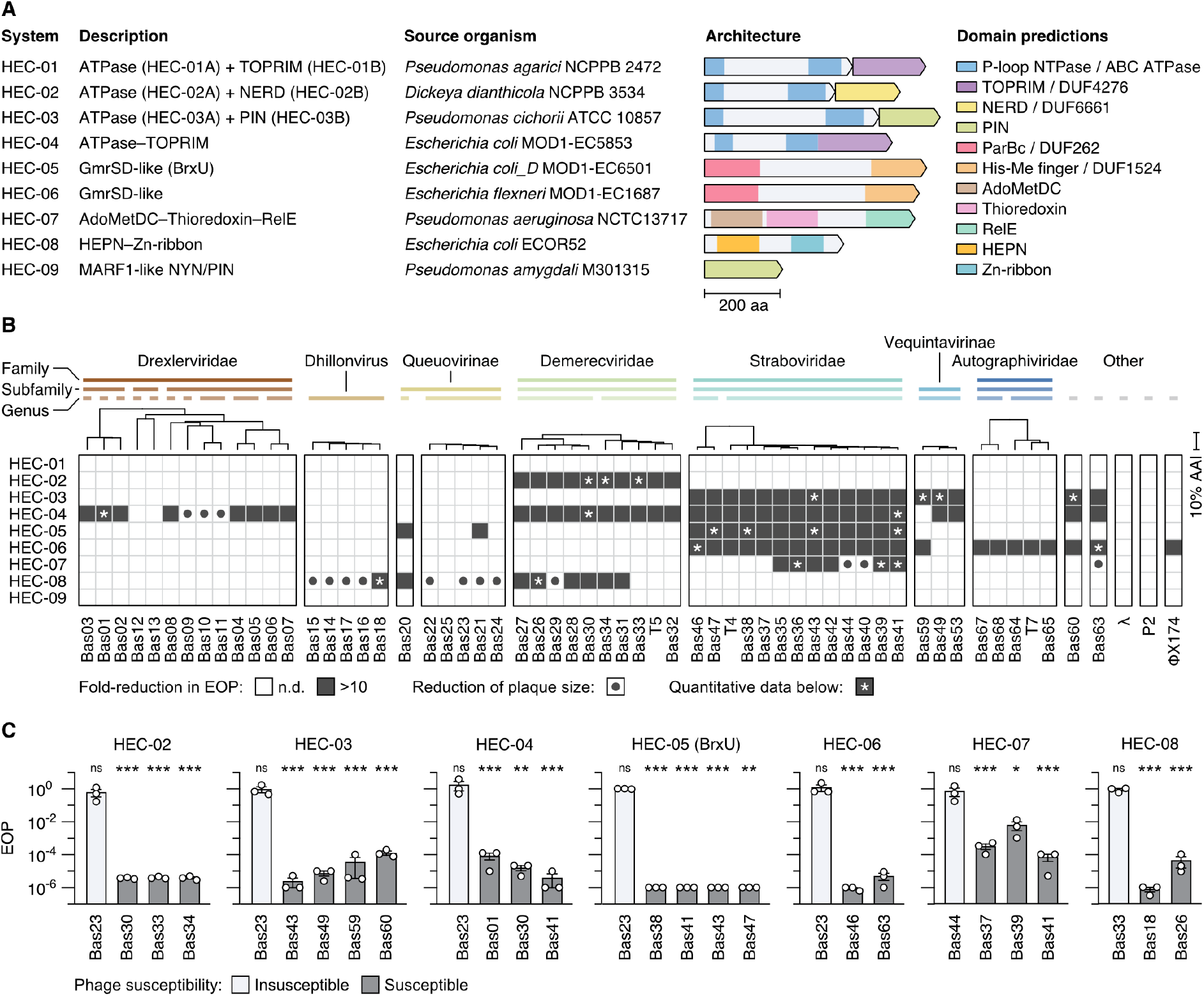
Hma-embedded candidate systems exhibit antiviral activity. (A) Candidate systems selected for testing. Domain predictions are based on Pfam annotations and structural comparison. ATPase: adenosine triphosphatase; TOPRIM: topoisomerase-primase; NERD: nuclease-related domain; PIN: PilT N-terminal; AdoMetDC: adenosylmethionine decarboxylase; HEPN: higher eukaryotes and prokaryotes nucleotide-binding; Zn-ribbon: Zinc ribbon; MARF1: meiosis regulator and mRNA stability factor 1; NYN: Nedd4-BP1/YacP nuclease. NTPase: nucleotide triphosphatase; ABC: ATP-binding cassette; DUF: domain of unknown function; His-Me: Histidine-metal. (B) Phage defence activity observed using a qualitative infection screen. Reduction in efficiency of plaquing (EOP) and plaque size represents the consensus of three replicates; n.d. = no reduction detected. Tree scale represents the percentage difference in average amino acid identity (AAI) between phage proteomes. Horizontal bars indicate phages belonging to the same taxa, grouped to the highest order taxonomic rank available (up to family). (C) Efficiency of plaquing (EOP) for select phages in the presence versus absence of HEC systems; *n* = 3, error bars represent the standard error of the mean. EOP for phages not susceptible to each respective HEC (i.e. Bas23, Bas33, and Bas44) are shown with white bars for comparison. Sample means were compared by *t*-test (Student, 1908) (comparing the plaque forming units (PFU)/mL of each phage in the presence of a HEC versus the non-defence control) with Holm-Šídák correction for multiple testing (Holm, 1979); ns = *p* > 0.05; * = *p* < 0.05; ** = *p* < 0.01; *** = *p <* 0.001. See **Figure S2** for assay results plotted by PFU/mL.

### Widespread phage defences with ABC ATPase domains

Based on sequence comparison, the ATPase domains of HEC-01 through HEC-04 appeared to be of the ATPases associated with diverse cellular activity (AAA/AAA+) clade. However, structure predictions revealed that these were instead ATP-binding cassette (ABC) ATPases. We identified similar ABC ATPase domains in several other defence systems, including PrrC, RloC, OLD, Gabija, Septu, Lamassu, Wadjet, Dnd, Pbe, PARIS, Ppl, PD-T4-4 and Menshen, some of which were previously misclassified as AAA/AAA+ ATPases (**Figure S4**). The distinguishing structural features of ABC ATPases have been described in detail elsewhere (Krishnan et al., 2020). In short, their core fold comprises a β sheet of five parallel strands flanked by three helices on one side and a single helix on the other, forming an αβα sandwich (Krishnan et al., 2020) (**Figure 3A–C**). In addition to this core, ABC ATPases are diversified by domain fusions to the N- or C-termini, or insertions between the first helix (H1) and second strand (S2) (Insert 1) or S2 and H2 (Insert 2) (Krishnan et al., 2020) (**Figure 3A–C**). In HEC-01A, HEC-02A, HEC-03A and HEC-04, the hairpin formed by the first preceding strand (PS1) and PS2 is extended into a β sheet by Insert 1 (**Figure 3D**), forming the characteristic open β-barrel-like structure reported to only be present in the ABC clade of P-loop nucleotide triphosphatases (NTPases) (Krishnan et al., 2020). Insert 1, Insert 2, and the C-terminal extensions differ substantially between the HEC ATPases (**Figure 3D**), consistent with their differences in phage defence specificity and diverse associated proteins.

**Figure 3.**
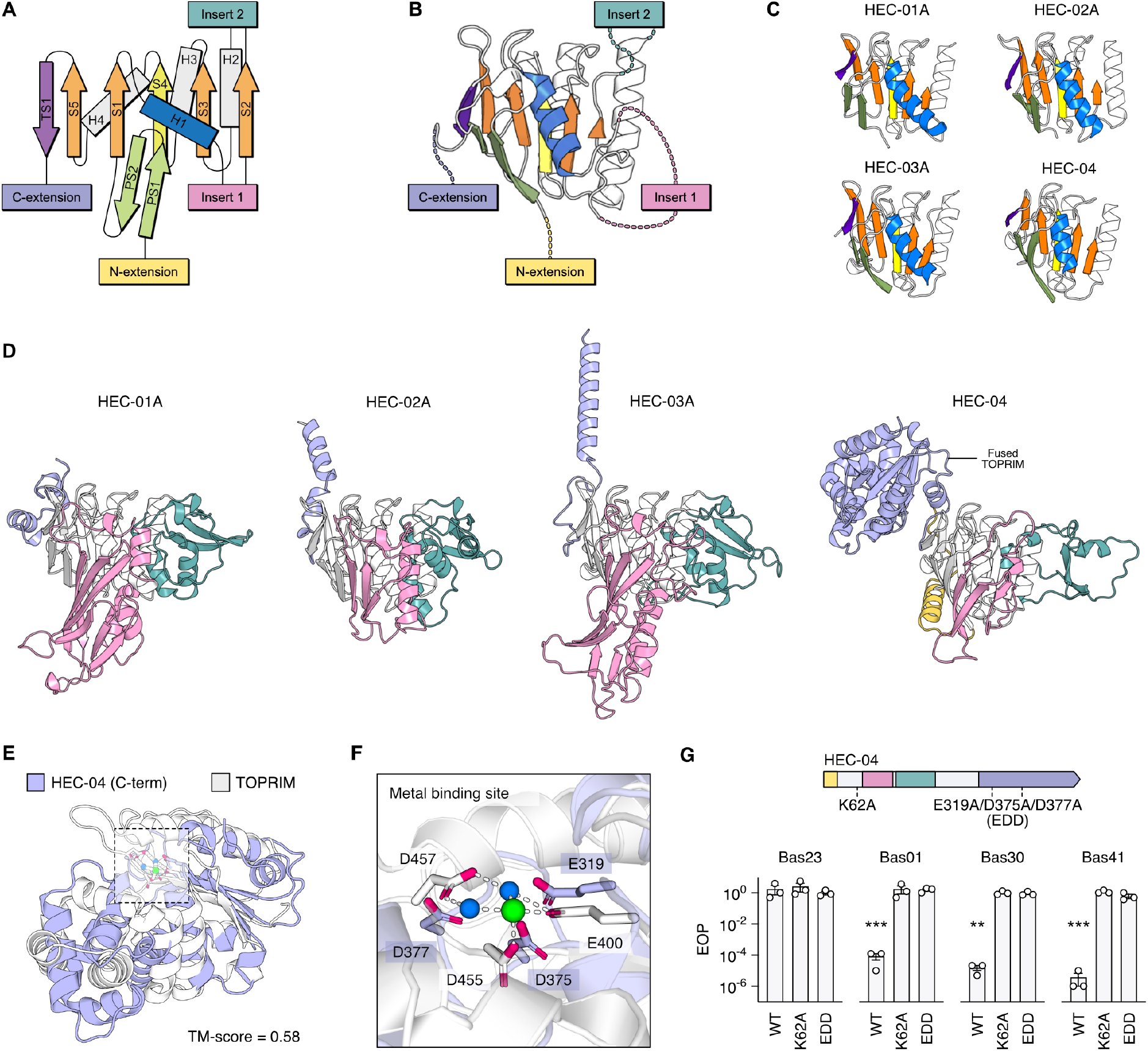
ABC ATPases are a common feature of phage defence systems. (A) Topology of the ABC ATPase core, adapted from (Krishnan et al., 2020). S; strand. H; helix. PS; preceding strand. TS; terminal strand. (B) ABC ATPase core of the structural maintenance of chromosomes (SMC) N-terminal head domain (PDB: 4I99). (C) Predicted structures of the ABC ATPase cores of HEC-01 through HEC-04. (D) Predicted structures of the complete ABC ATPase proteins of HEC-01 through HEC-04. (E) Structural alignment of the TOPRIM domain from the predicted structure of HEC-04 (purple) and *Burkholderia pseudomallei* (Bp) OLD (PDB: 6NK8; white). Template modelling (TM)-score was calculated with US-align (Zhang et al., 2022), where TM-score ≥ 0.5 suggests shared global topology (Xu and Zhang, 2010). The metal binding site presented in panel E is highlighted. (F). The metal binding site of Bp OLD and corresponding residues of HEC-04. Water molecules are shown in blue and the metal ion is shown in green. Relevant hydrogen bonds between Bp OLD residues, water atoms and the metal ion are represented by white dashed lines. (G) EOP data for HEC-04; Wild type (WT), Walker A mutant (K62A), and metal-binding site mutant (EDD; E319A, D375A, D377A); *n* = 3, error bars represent the standard error of the mean. Sample means were compared by *t*-test (Student, 1908) (comparing the PFU/mL of each phage in the presence of a WT or mutated HEC versus the non-defence control) with Holm-Šídák correction for multiple testing (Holm, 1979); ** = *p* < 0.01; *** = *p* < 0.001. See **Figure S2** for assay results plotted by PFU/mL. See **Figure S5** for alignment coverage, predicted local distance difference test (pLDDT), and predicted aligned error (PAE) plots associated with each HEC structural prediction.

Aside from the differences between their ATPases, the additional proteins (or additional domain for HEC-04) associated with HEC-01 through HEC-04 vary. HEC-01B and the C-terminal extension of HEC-04 share structural similarity with TOPRIM domain-containing proteins. ABC ATPase–TOPRIM associations are found in several other systems, including overcoming lysogenisation defect (OLD) nucleases, Gabija, PARIS, and Wadjet systems, though the ATPase components of all these systems are structurally distinct (**Figure S4A**). When aligned to the TOPRIM domain of *Burkholderia pseudomallei* (Bp) OLD, HEC-01 and HEC-04 share a conserved metal binding site important for nuclease activity (Schiltz et al., 2019) (**Figure S6A,B; Figure 3E,F**). For HEC-04, mutation of key active site residues in either the ABC ATPase or nuclease domains demonstrated their essential roles in defence (**Figure 3G**). However, no defence was detected for HEC-01 in our assays (**Figure 2A**), potentially due to incompatibility with the heterologous host or specificity outside the scope of phages tested. HEC-02B is similar to NERD proteins (**Figure S6C**), a class of nucleases belonging to the PD-(D/E)xK phosphodiesterase superfamily (Grynberg, 2004; Steczkiewicz et al., 2012), and contains the signature motif (D/E/Q)xK associated with phosphodiesterase activity (Aravind, 2000b; Grynberg, 2004; Knizewski et al., 2007) (**Figure S6D**). NERD proteins were previously implicated in phage defence based on contextual analysis (Makarova et al., 2011; Bernheim et al., 2021), which we have confirmed here experimentally (**Figure S3A**). HEC-03B contains a PIN domain similar to those of VapC family ribonucleases found in VapBC toxin-antitoxin systems (**Figure S6E,F**). However, the HEC-03A ABC ATPase bears no relation to VapB antitoxins. Overall, our findings expand the large superfamily of phage defence systems comprised of ABC ATPase domain-containing proteins coupled with diverse effectors (Krishnan et al., 2020).

### A family of GmrSD-like phage defences

Genes encoding GmrSD-like proteins were also frequently embedded in Hma systems, from which we selected HEC-05 and HEC-06 as candidates. GmrSD is a type IV restriction-modification (RM) system, initially reported as a two-gene system in *E. coli* CT596 (EcoCTGmrSD) encoding separate DUF262 (GmrS) and DUF1524 (GmrD) domain-containing proteins (Bair and Black, 2007). However, EcoCTGmrSD was later found to be a single multi-domain protein where the DUF262 and DUF1524 domains were joined by a large linker domain, and this architecture was found to be typical for GmrSD homologs (Machnicka et al., 2015). Two other types of GmrSD-like proteins are known to have roles in phage defence: the phosphorothioation-dependent nicking endonuclease SspE (Xiong et al., 2020), and the BREX-associated BrxU (Picton et al., 2021; Mariano and Blower, 2023) (of which the chosen HEC-05 homolog shares 99.2% amino acid identity). However, HEC-06 shares only 23.7% amino acid identity across 11% alignment coverage with BrxU. Structures for BrxU and SspE have been solved experimentally, both forming intertwined homodimers with distinct quaternary structures (Xiong et al., 2020; Picton et al., 2021). AlphaFold accurately reproduced the individual domains of BrxU and SspE, but did not replicate the relative position of the DUF262 domain (**Figure S7A,B**). Aligning the predicted structure of HEC-06 with either the known structure of BrxU or SspE required the introduction of multiple twists outside the flexible linker region, suggesting that the overall quaternary structure of a HEC-06 complex is distinct from these other GmrSD-like proteins (**Figure S7C,D**). Additionally, comparison between the individual domains of HEC-06, GmrSD, BrxU and SspE also suggests that HEC-06 is distinct from other GmrSD-like proteins (**Figure 4A,B**). These variations in structure are consistent with the differences in phage defence we observed for HEC-06 versus HEC-05 (BrxU) (**Figure 2B**).

**Figure 4.**
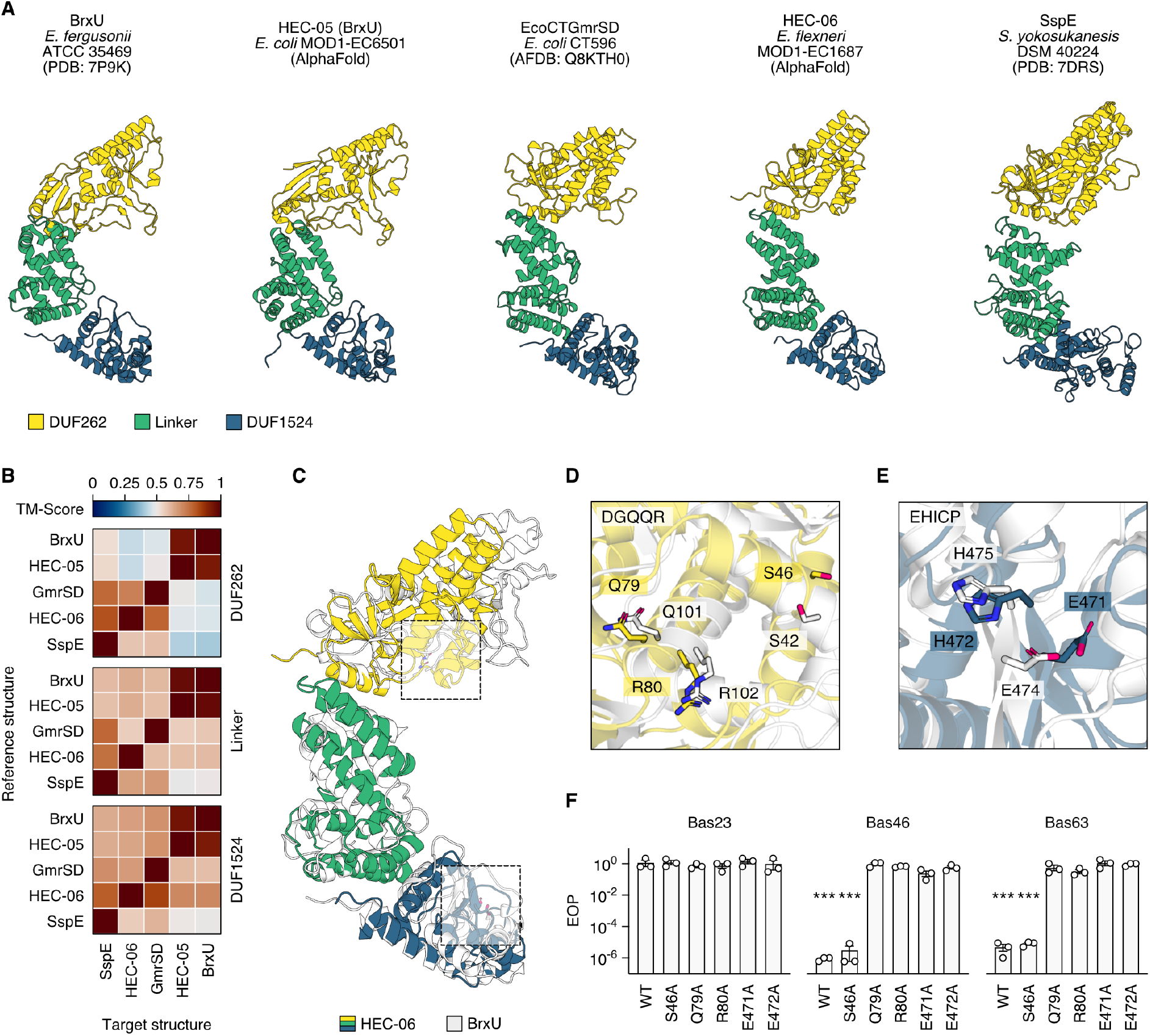
Diverse GmrSD-like proteins provide antiviral defence. (A) Structures of five GmrSD-like proteins. Each domain was aligned independently for better comparability between proteins. These ‘straightened’ representations do not reflect the expected bent conformation found in experimentally derived structures where flexibility between each of the domains allows for the formation of intertwined dimers (**Figure S7A,B**). AFDB: AlphaFold database. (B) Pairwise TM-scores calculated between each domain of the GmrSD-like proteins. Scores are normalised by the length of each reference structure. (C) Structural alignment of the predicted HEC-06 structure (coloured) with BrxU (PDB: 7P9K) (white). Each domain was aligned independently as above. Sites highlighted by boxes are enlarged in panels D and E. (D,E) Residues targeted for S46A and nucleotide binding/hydrolysis mutants (Q79A, R80A) (D) and nuclease motif mutants (E471A, H472A) (E), aligned with the corresponding residues of BrxU. (F) Efficiency of plaquing data for WT and mutant HEC-06; *n* = 3, error bars represent the standard error of the mean. Sample means were compared by *t*-test (Student, 1908) (comparing the PFU/mL of each phage in the presence of a WT or mutated HEC versus the non-defence control) with Holm-Šídák correction for multiple testing (Holm, 1979); *** = *p* < 0.001. See **Figure S2** for assay results plotted by PFU/mL.

HEC-06, EcoCTGmrSD, and SspE do not contain the predicted nucleotide-binding (RLFDS) motif present in the DUF262 domain of BrxU (Machnicka et al., 2015; Picton et al., 2021). However, HEC-06 contains a serine (S46) that protrudes from the second α-helix of the DUF262 domain (and is conserved in SspE) that may be analogous to S42 of BrxU (**Figure 4C,D**). In BrxU, mutation of this residue prevents dimerization, which is essential for phage defence (Picton et al., 2021). However, mutation of the serine (S46A) in HEC-06 did not impair the observed antiphage activity (**Figure 4F**). The DUF262 domain of HEC-06 also contains the DGQQR motif conserved in other GmrSD-like proteins, which is involved in nucleotide hydrolysis (Machnicka et al., 2015; Picton et al., 2021) (**Figure 4C,D**). Accordingly, Q79A and R80A mutations in this motif resulted in loss of phage resistance (**Figure 4F**). The DUF1524 domain of HEC-06 contains a predicted HNH nuclease motif (EHICP) that aligns with that in BrxU (DHIYP) (**Figure 4C,E**). Mutation of the acidic residue (aspartic acid) and conserved histidine to alanine in BrxU eliminated phage resistance (Picton et al., 2021), as did mutation of the glutamic acid (E471A) and histidine (H472A) in HEC-06 (**Figure 4F**). Overall, our findings expand the known family of GmrSD-like defence systems and suggest further research is warranted to understand the basis for co-option of this widespread family for phage defence.

### Diverse RNase-based defences

Finally, we identified several diverse families of putative RNases (HEC-07 through HEC-09). The N-terminal domain of HEC-07 shares structural similarity with the adenosylmethionine decarboxylase (AdoMetDC)-like domain of spermine synthase (**Figure 5A**). The second domain is similar to thioredoxin but lacks the conserved thioredoxin Cx2C active site (Ingles-Prieto et al., 2013) (**Figure 5A**). The C-terminal domain is similar to the *Helicobacter pylori* RelE family RNase toxin HP0892 (Pathak et al., 2013) (**Figure 5A**) and contains a conserved histidine (H543) known to be important for the RNase activity of several other RelE family toxins (Pathak et al., 2013). HEC-08 comprises an N-terminal DUF3644 domain, previously characterised as a HEPN domain (Anantharaman et al., 2013), and several C-terminal zinc-finger-like domains (**Figure 5B**). Comparison of the N-terminal domain with PDB structures revealed matches to other HEPN domain-containing proteins, including *Shewanella oneidensis* toxin SO_3166 (**Figure 5B**). Mutation of the histidine of the Rx4-6H motif (H128) conserved in HEC-08 and other HEPN domain-containing proteins negated defence (**Figure 5C,D**). The predicted C-terminal structure of HEC-08 had no similarity to known structures, but had sequence similarity to a zinc beta ribbon. Lastly, HEC-09 had weak N-terminal sequence similarity to an initiation factor 2B/5 domain (helix-turn-helix protein) and a C-terminal PIN-like domain. Structural comparison revealed similarity to the NYN domains of human and mouse meiosis regulator and mRNA stability factor 1 (MARF1) proteins (**Figure 5E**). HEC-09 shares three conserved aspartates with human MARF1 (D358, D426 and D452) that form a negatively charged pocket known to be the active site for metal binding and RNase activity. However, HEC-09 did not provide defence against any of the phages we tested.

**Figure 5.**
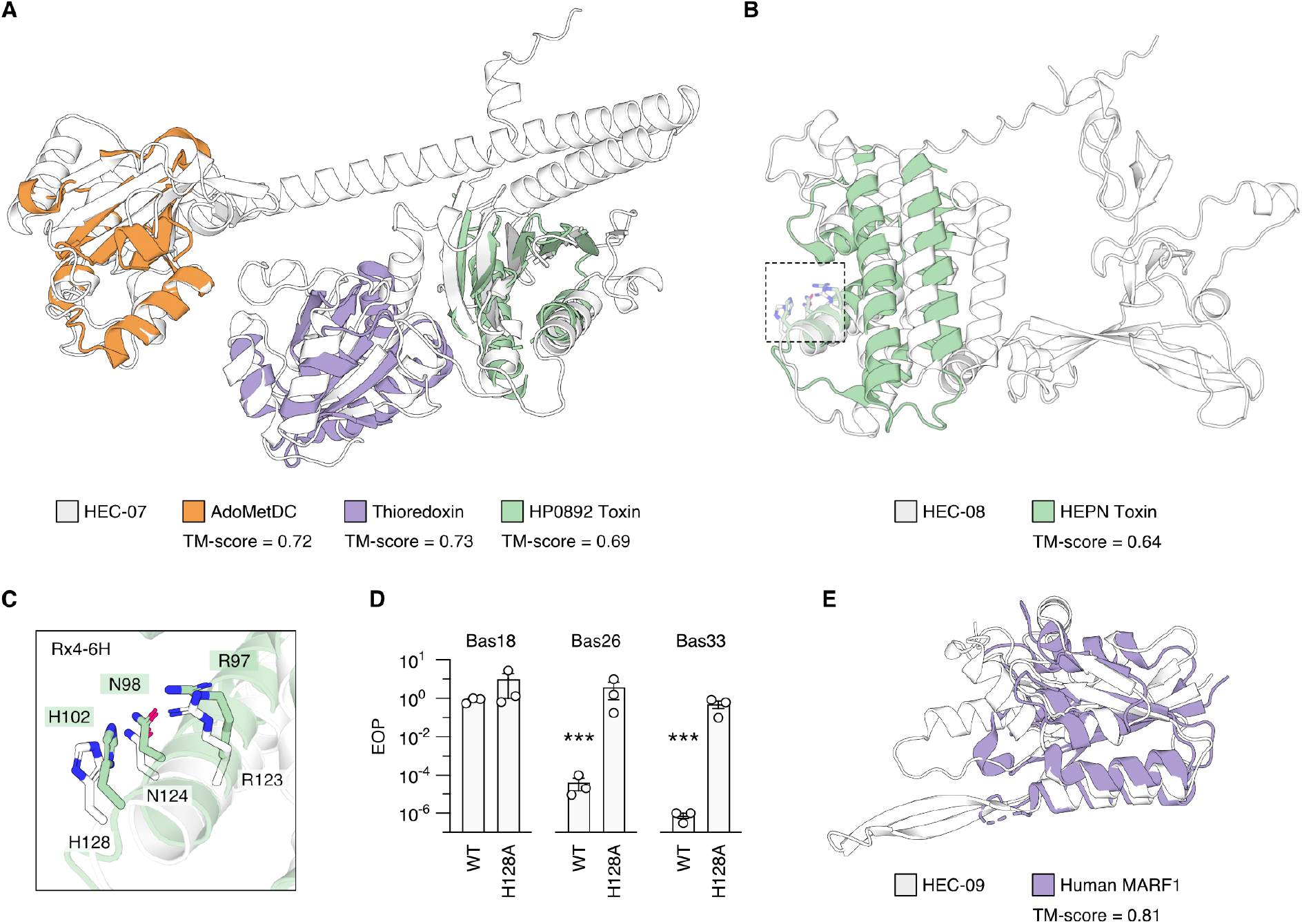
Putative RNase-based defence systems. (A) The structures of the AdoMetDC-like domain of spermine synthase (PDB: 3C6K; orange), a thioredoxin (PDB: 3VFI; purple) and the *Helicobacter pylori* toxin HP0892 (PDB: 4LTT; green), aligned to the predicted structure of HEC-07 (white). (B) The structure of *Shewanella oneidensis* HEPN-domain containing toxin SO_3166 (PDB: 5YEP; green) aligned to the predicted structure of HEC-08 (white). The Rx4-6H motif presented in panel C is highlighted. (C) The Rx4-6H motif conserved between the HEPN domain-containing toxin and HEC-08, targeted for mutagenesis. (D) Efficiency of plaquing (EOP) data from mutational analysis of HEC-08; *n* = 3, error bars represent the standard error of the mean. Sample means were compared by *t*-test (Student, 1908) (comparing the PFU/mL of each phage in the presence of a WT or mutated HEC versus the non-defence control) with Holm-Šídák correction for multiple testing (Holm, 1979); *** = *p* < 0.001. See **Figure S2** for assay results plotted by PFU/mL. (E) The structure of Human MARF1 (PDB: 6FDL; purple) aligned to the predicted structure of HEC-09 (white).

### Guilt-by-embedding implicates hundreds of putative new defence systems

Based on our success predicting functional defence families embedded in Hma systems, we hypothesised that additional defence candidates could be found embedded in other types of systems. In particular, BREX, DISARM, and Type I RM systems also frequently contained embedded genes (**Figure S8A,B**). Using a similar approach as described for Hma, we identified 17,432 nonredundant BREX/DISARM/RM-embedded genes, encoding 3,231 protein families, which were observed as 3,568 unique combinations of gene cassettes (**Figure 6A**). We searched for additional homologs of these cassettes (including non-embedded) in RefSeq v209 genomes, then curated this list of candidate systems to exclude those likely to encode known defence systems, MAPs, or proteins with non–defence associated functions, to derive a set of 145 single- and multi-gene <u>p</u>hage <u>d</u>efence <u>c</u>andidate (PDC) systems (**Figure 6A; Table S3-8**). To enable future functional studies of these PDC systems, we have added them to our bioinformatic defence system identification tool, PADLOC, and associated webserver (https://padloc.otago.ac.nz).

**Figure 6.**
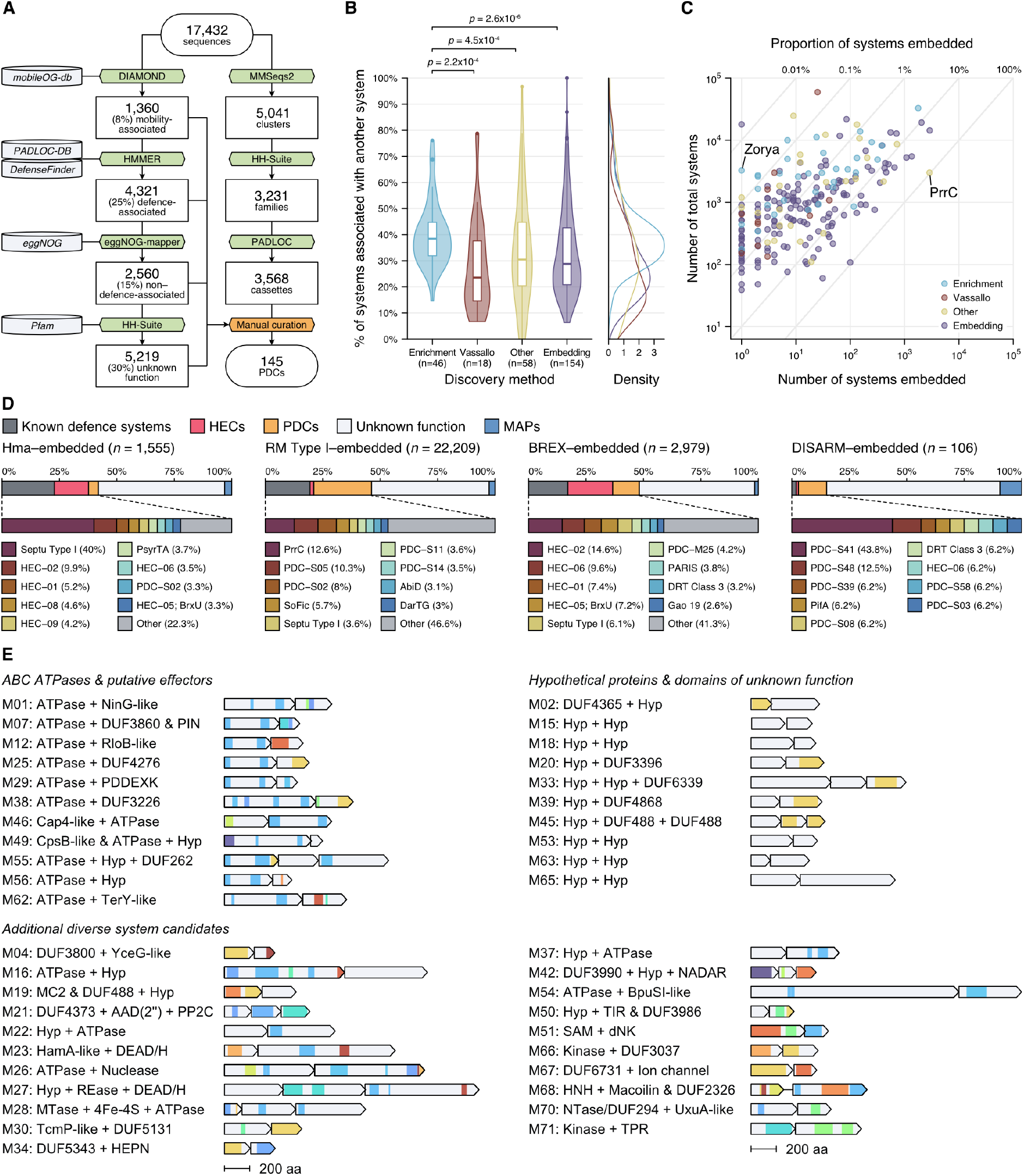
New phage defence candidates can be predicted through guilt-by-embedding. (A) Flow chart of embedded protein function assignment. Tools are highlighted in green, and their associated databases in grey. Filtering steps are highlighted in orange. (B) Proportion of all defence system families associated with other defence systems (genetically separated by < 5 ORFs), grouped by method of discovery. The ‘Enrichment’ category includes systems discovered through defence enrichment scoring algorithms, from Doron *et al*., 2018 (Doron et al., 2018), Gao *et al*., 2020 (Gao et al., 2020), and Millman *et al*., 2022 (Millman et al., 2022). The ‘Vassallo’ category includes systems discovered through the functional screen by Vassallo *et al*., (Vassallo et al., 2022). The ‘Other’ systems include all other systems included in the PADLOC database (excluding those identified herein). Group distributions were compared with a Kruskal-Wallis rank sum test (Kruskal and Wallis, 1952): χ^2^ = 13.42; *p* = 3.8x10^−3^, followed by a *post hoc* Wilcoxon rank sum test (Wilcoxon, 1945) with Benjamini-Hochberg correction for multiple testing (Benjamini and Hochberg, 1995). Comparisons with *p* < 0.05 are shown. (C) Proportion of systems embedded in BREX, DISARM, and RM Type I systems. Each point represents a different type of system. Zorya type II is labelled as an example of a widespread system for which we identified just a single embedded instance. PrrC is labelled as an example of a system almost exclusively found embedded in type I RM systems. (D) Proportion of embedded genes identified as encoding known defence systems (grey, further detailed underneath), HECs (red), PDCs (orange), MAPs (blue), or proteins with otherwise unknown functions (white). (E) Examples of some multi-gene PDCs. Hyp: hypothetical protein; MC2: ABC-3C middle component 2; AAD(2’’): Aminoglycoside-2’’-adenylyltransferase; PP2C: protein phosphatase 2C; DEAD/H: DEAD/H-box helicase; 4Fe-4S: 4Fe-4S iron-sulfur cluster binding; NADAR: NAD and ADP-ribose; TIR: toll/interleukin-1 receptor; SAM: radical S-adenosyl-L-methionine; dNK: deoxyribonucleoside kinase; TPR: tetracopeptide repeat.

Despite the success of defence island enrichment–based approaches in discovering new defence systems, genetic co-localisation of systems is less common than ‘solo’ systems and high-throughput functional screening demonstrates many systems likely await discovery (Vassallo et al., 2022). Overall, the systems we predicted through embedding (HECs and PDCs) have a significantly lower association with other defence systems than those identified by defence enrichment analyses (**Figure 6B**). We propose that a single observed embedded instance is sufficient to warrant further investigation into potential phage defence function (after excluding mobilome- and other non–defence associated functions). For example, we found single instances of embedded Aditi, Bunzi, PD-T4-5, DRT type III, and Zorya type II systems versus 231, 585, 1,188, 1,273, and 1,590 non-embedded homologs, respectively. Many other types of systems are found embedded only a few times (**Figure 6C**). With the addition of our top 145 PDCs, more than 50% of the genes embedded in each of Hma, BREX, DISARM, and RM Type I do not have a clear mobility-, defence-, or non–defence associated function, and likely contain additional undiscovered defence systems that are rare (below our arbitrary abundance cut-offs) in the genomes searched (**Figure 6D; Figure S8C**). Together, these results demonstrate that most genes embedded in defence systems are not MGEs or genes with metabolic or housekeeping functions, instead they likely constitute a wealth of new phage defence systems (**Figure 6E**).

## DISCUSSION

Efforts to identify genes overrepresented in the genetic context of known defence systems have greatly expanded our view of the prokaryotic pan-immune system (Doron et al., 2018; Gao et al., 2020; Millman et al., 2022). However, a recent experimental screen revealed multiple defence systems not identified via earlier bioinformatic approaches, highlighting the likely vast repertoire of phage defence systems awaiting discovery (Vassallo et al., 2022). Here, we demonstrated guilt-by-embedding as an alternative approach to system discovery— investigating genes that are embedded within known defence systems without relying on their specific enrichment with other systems. Overall, we predicted more than 150 new types of phage defence systems, confirmed the activity of six new types and the earlier independent discovery of the BrxU system (Picton et al., 2021). We have made the 145 untested phage defence candidates searchable with our bioinformatics tool PADLOC (Payne et al., 2021, 2022).

The most common gene family we identified embedded in Hma systems encoded many diverse ABC ATPases, which were associated with additional genes encoding TOPRIM, NERD, or PIN domain-containing proteins (HEC-01, HEC-02 and HEC-03, respectively). ABC ATPases are found in many defence systems, including PrrC, RloC, OLD family nucleases, Gabija (GajA), Septu (PtuA), Lamassu (LmuB), Wadjet (JetC), phosphorothioation-based restriction systems (DndD, PbeC), PARIS (AriA), Ppl, PD-T4-4 and Menshen (MenB) (Davidov and Kaufmann, 2008; Xu et al., 2010; Anantharaman et al., 2013; Doron et al., 2018; Schiltz et al., 2019; Xiong et al., 2019; Gao et al., 2020; Millman et al., 2022; Rousset et al., 2022). Through extensive remodelling of the first and second inserts of the ABC ATPase, N- and C-terminal fusions, and co-evolution with different effectors, ABC ATPase–based defence systems may have evolved as diversified phage sensors that activate a fused effector domain (e.g. the TOPRIM domain of HEC-04) or associated effector proteins (e.g. the NERD or PIN domain-containing proteins of HEC-02 and HEC-03, respectively) to elicit defence. How these HEC-associated ATPases sense infection and whether it is ATP binding or hydrolysis which is important for function remains to be determined. At least for HEC-04, a functioning Walker-A motif is necessary for defence. The specificity of ABC ATPase–based defence systems likely depends on the variations found in the first and second inserts, which may alter the sensitivity of the ATPases to different triggers. Several known triggers of these systems are phage-encoded anti-defence proteins. For example, the DNA mimic Ocr activates PARIS (Rousset et al., 2022) and Gabija systems (Stokar-Avihail et al., 2023), the RM inhibitor Stp activates PARIS (Rousset et al., 2022) and PrrC systems (Penner et al., 1995), and RecBCD inhibitors activate OLD nucleases (Lindahl et al., 1970; Myung and Calendar, 1995). Some ABC ATPases, such as LmuB, associate with many different effectors, such as nucleases, proteases, and sirtuins (Krishnan et al., 2020; Millman et al., 2022), while others are more discriminative. Likewise, the same type of effector may be associated with several different ABC ATPases. For example, TOPRIM domains are found paired with OLD, Gabija, PARIS, and Wadjet ABC ATPases, and may have evolved alongside these ABC ATPases as indiscriminate nucleic acid degradation systems that induce dormancy or death as a second-line defence strategy (Krishnan et al., 2020). Our developing understanding of several recurring themes in defence systems, such as the extensive diversification of many ABC ATPase-based defences, demonstrates the importance of exploring how ‘novel’ defence systems fit into the larger evolutionary history of prokaryotic immunity.

The biological significance of defence system embedding versus the formation of islands of neighbouring systems or systems co-residing in hosts at different genetic loci remains to be explored. Embedding may occur as an outcome of the ‘genomic sink’ hypothesis (Makarova et al., 2011), where defence system loci are more amenable to the insertion of new genes than other parts of the genome. Once a system becomes embedded, a selfish benefit may be conferred to the embedded system under selective pressure for the surrounding system, as horizontal transfer of the systems would be linked. Additionally, embedding may provide functional advantages that benefit the host. Some embedded systems provide backup mechanisms of defence upon inactivation of the surrounding system. For example, the anticodon nuclease PrrC, embedded in the *Eco*prrI (*hsdMSR*) restriction modification system, is kept catalytically inactive by the restriction modification system unless *Eco*prrI is inactivated by the Stp anti-restriction protein encoded by T4-like phages (Meineke and Shuman, 2012). The embedding of other ABC ATPase–based defences, which are triggered by different anti-defence proteins, may also be beneficial. For example, a phage-encoded Ocr may inhibit BREX (Isaev et al., 2020), but trigger an embedded PARIS or Gabija system. Similarly, some embedded systems provide redundancy or activity to counteract phage evasion of the primary system. For instance, the BREX system encoded on the plasmid pEFER is assisted by the embedded system BrxU through contrasting defence strategies of targeting unmodified and modified DNA, respectively (Picton et al., 2021). Here, we find that BrxU and other GmrSD-like proteins are also frequently embedded in Hma and RM Type I systems, which may perform a similar role in these contexts. System embedding might also enable efficient co-regulation of defences, as genes embedded in Hma, BREX, DISARM, and RM Type I systems are in the same orientation as the flanking system in 96.7% of cases.

We demonstrated that most genes embedded in the defence systems we investigated encode known systems, new systems, and other diverse proteins with domains associated with defence, or with little similarity to known sequence families. The embedded genes that do not map to known defences constitute a wealth of untapped candidates for the discovery of new defence systems. Exploring the candidates we have highlighted here, and the many genes embedded in other defence systems, will be valuable for advancing our understanding of prokaryotic antiviral responses. Expanding our knowledge of the immune arsenal of prokaryotes has important implications for protecting industrial bioproduction strains, developing phage therapies against bacterial pathogens, and discovering new enzymatic functions to expand the molecular biology and biotechnology toolkit.

## MATERIAL AND METHODS

### Identification of Hma-embedded genes and Hma-embedded candidate selection

PADLOC v1.1.0 (Payne et al., 2021, 2022) with PADLOC-DB v1.4.0 was used to search for Hma systems in all RefSeq v209 Archaea and Bacteria genomes (*n* = 234,099), allowing up to three embedded genes between each *hma* gene. This threshold was limited to three genes to avoid spurious identification of Hma systems. The identified Hma systems were curated to include only high-quality predictions where each expected *hma* gene occurred only once, all *hma* genes were on the same strand, and none were pseudogenes. The protein sequences of embedded genes were retrieved and duplicate sequences removed with seqkit v2.3.0 (Shen et al., 2016). Any complete known defence systems were also removed. The remaining proteins were grouped into clusters at 30% minimum sequence ID and 80% minimum alignment coverage using MMseqs2 v14.7e284 (Steinegger and Söding, 2017, 2018). Clusters were further grouped into families based on an all-against-all HMM comparison with HHsearch v3.3.0 (Steinegger et al., 2019) with cut-offs of 70% minimum probability and 70% alignment coverage for both target and query. For gene cassettes embedded in more than five Hma systems, HMMs were built from each protein cluster using HMMER v3.3.2 (Eddy, 2011). These gene cassettes were searched back across all of the RefSeq genomes using PADLOC and filtered for those present in more than 100 genomes. To generate **Figure S1**, phyla were grouped using GTDB v207 (Parks et al., 2022) taxonomy.

### Protein structure prediction and comparison

Structure prediction was performed with ColabFold v1.2.0 (Jumper et al., 2021; Mirdita et al., 2022). See **Figure S5** for alignment coverage, predicted local distance difference test (pLDDT), and predicted aligned error (PAE) plots associated with each HEC structural prediction. Structural similarity searches were performed using the DALI (Holm, 2020), Foldseek (van Kempen et al., 2022) and FATCAT (Li et al., 2020) web servers. Structural alignments were guided by DALI, FATCAT, US-align v20230609 (Zhang et al., 2022) and the PyMOL superposition algorithm (Schrödinger, LLC, 2021). Structural images were generated using PyMOL.

### Cloning of candidate defence systems

Sequence and source details for cloned systems are listed in **Table S2**. Each Hma-embedded candidate was commercially synthesised, cloned and sequence-validated by Twist Bioscience. The candidates were cloned into the pTwist Chlor Medium Copy vector, which contains a p15A origin of replication and chloramphenicol resistance marker. Where possible, the putative native promoter of each locus was included (∼150 bp upstream), in addition to an IPTG-inducible T5/lac promoter. *E. coli* K-12 MG1655 ΔRM was transformed by heat shock.

### Phage plaque assays

*Escherichia coli* BASEL phages, P2vir, lambdavir and T4 were obtained from the BASEL phage collection (University of Basel, Switzerland). Phages T5, T7 and phiX174 were obtained from Chase Beisel (Helmholtz Centre for Infection Research, Germany). For all plaque assays, *E. coli* K-12 MG1655 ΔRM containing the pTwist vector with a candidate defence system or non-defence control (encoding mCherry) were grown in 6 mL Lysogeny Broth (LB) supplemented with 25 μg mL^-1^ chloramphenicol overnight at 37°C with shaking at 200 rpm. A 300 μL aliquot of the overnight cultures were mixed with 8mL of 0.35% w/v LB agar supplemented with 20 mM of MgSO_4_ and 5 mM of CaCl_2_ and poured over 1.5% w/v LB agar supplemented with 25 μg mL^-1^ chloramphenicol and 0.1 mM IPTG (strains expressing HEC-03 and HEC-04 were not supplemented with IPTG due to apparent toxicity). Phages were spotted onto the bacterial overlays and incubated overnight at 30°C until plaques could be counted. For the initial qualitative assessment of plaque reduction, 8 μL of a 10-fold dilution series of phage were first spotted onto *E. coli* K-12 MG1655 ΔRM to determine the most dilute concentration where >10 plaques could be observed. From this dilution, 8 μL of each phage were spotted onto the bacterial overlays and the efficiency of plaquing was determined by comparing plaque formation in bacteria containing the defence systems with those containing the mCherry control. For quantitative efficiency of plaquing assays, 10-fold dilution series of phage were plated onto the bacterial overlay. The efficiency of plaquing (EOP) was determined as above. AAI and hierarchical clustering of phage genomes was calculated using EzAAI (Kim et al., 2021).

### Identification of BREX–, DISARM– and RM Type I–embedded genes and phage defence candidate selection

BREX, DISARM and RM Type I systems were filtered for those with embedded genes, where all system genes were encoded on the same strand, and no genes were pseudogenes. The proteins encoded up to three ORFs either side of each system were clustered with MMseqs2 (30% minimum sequence ID and 80% minimum coverage). One representative system was selected per gene cluster based on this grouping. Proteins encoded by BREX, DISARM and RM Type I–embedded genes were compared against mobileOG-DB v1.6 (Brown et al., 2022) sequences using the *blastp* function of DIAMOND v2.1.8 (Buchfink et al., 2021) with default parameters. Putative hits were filtered for those with both query and target coverage >=70%. Comparisons with PADLOC-DB v1.4.0 (Payne et al., 2021) and DefenseFinder v1.2.3 (Tesson et al., 2022) HMMs were done using the *hmmsearch* function of HMMER v3.3.2 (Eddy, 2011) with default parameters. Putative hits were filtered for those with full sequence bitscores >=50. Comparison with eggNOG orthologous groups was done using the eggNOG-mapper v2.0.1 (Cantalapiedra et al., 2021) webserver with default parameters. Comparison with Pfam v35 (El-Gebali et al., 2019) was done using HHsearch v3.3.0 (Steinegger et al., 2019) with default parameters. Putative hits were filtered for those with probability >= 80%. Putative functions were assigned to the proteins encoded by embedded genes based on the following priority: mobility-associated proteins (MAPs) based on similarity to mobileOG-DB sequences, excluding proteins in the ‘element stability, transfer, or defense’ category; defence-associated proteins based on similarity to PADLOC-DB and DefenseFinder; non–defence-associated proteins based on similarity to eggNOG orthologous groups, excluding those in the ‘defense mechanisms’ (V), and ‘function unknown’ (S) functional categories. To generate the final PDC models for PADLOC, all PDC proteins identified in the entire RefSeq v209 database were clustered with MMseqs2, and an HMM was built from each cluster that contained at least one of the original sequences identified embedded in a BREX, DISARM or RM Type I system. For **Figure 6B**, defence systems were considered to be associated with another system if they had no more than five ORFs between them. Approximately 27.6% of the known systems identified in RefSeq v209 genomes did not have additional systems associated with them, but also had five or less ORFs between them and at least one contig boundary. To avoid such cases potentially skewing the data towards artificially lower association frequencies, any system separated from a contig boundary by five or less ORFs was excluded from this analysis.

## Supporting information

Supplemental Table Descriptions

## ACKNOWLEDGEMENT

We are grateful to Alexander Harms and his group for providing the BASEL phage collection (Maffei et al., 2021). We are grateful to Chase Beisel and his group, particularly Frank Englert, for providing T5, T7 and phiX174 phages. We thank members of the Phage Host Interactions laboratory (Otago) for helpful discussions. We acknowledge the use of the New Zealand eScience Infrastructure (NeSI) high-performance computing facilities in this research. The NeSI facilities are provided by and funded jointly by the NeSI collaborator institutions and through the Ministry of Business, Innovation and Employment’s Research Infrastructure programme.

