## Supplemental Figures for "New antiviral defences are genetically embedded within prokaryotic immune systems"

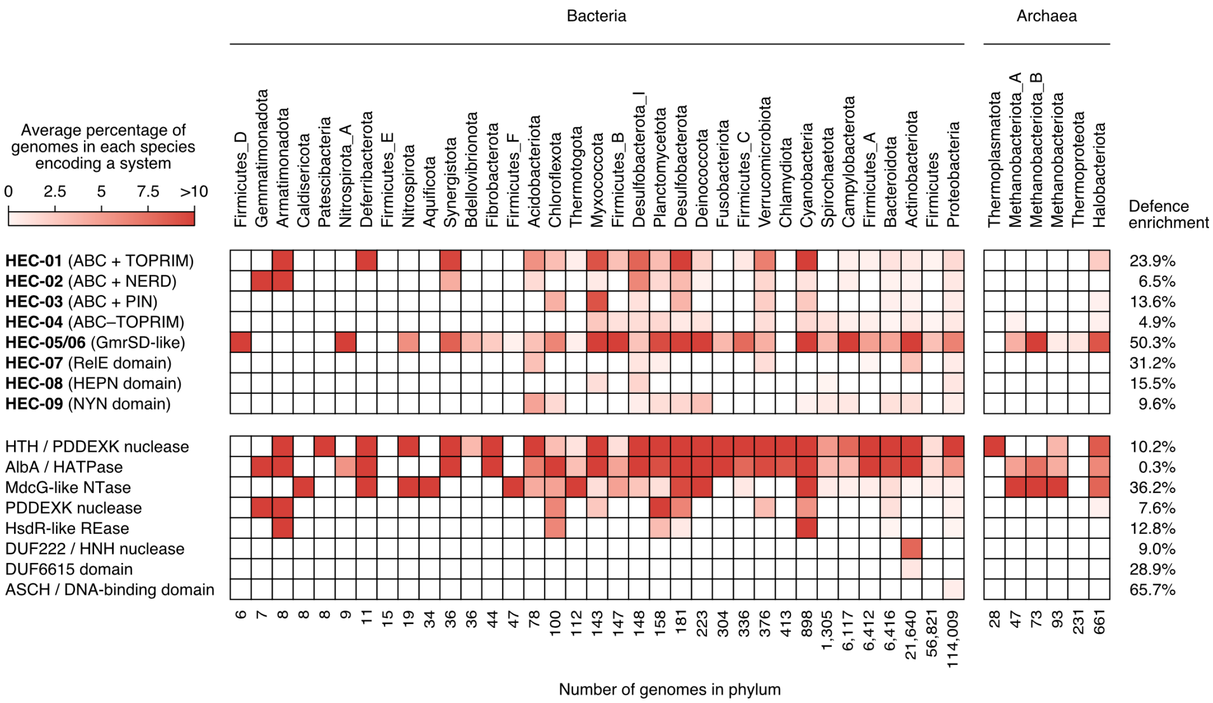


**Figure S1. Clusters of genes commonly embedded in Hma systems are also generally widespread.** The abundance of Hma-embedded gene cassettes identified in bacterial and archaeal genomes, irrespective of Hma association. Only cassettes identified in more than 100 genomes are shown. Only RefSeq v209 genomes which were also present in the GTDB v207 taxonomy database are shown (approximately 78% of all genomes searched). Phyla are grouped by GTDB v207 taxonomy. The heatmap represents, for each phylum, the average percentage of genomes in each species of that phylum encoding a system. System prevalence was weighted this way to limit biases in phyla that contain many closely related genomes of the same species. Putative systems selected for analysis were assigned unique HEC (Hma-embedded candidate) identifiers. Putative systems not selected for analysis are assigned general names based on their Pfam domain assignments (a slash '/' signifies multiple domains in the same protein). Defence enrichment is based on all genomes searched, where a candidate is classified as being associated with another defence system if they are separated by less than five open reading frames. HTH: Helix-turn-helix; HATPase: Histidine kinase–like ATPase; REase: Restriction endonuclease; NTase: Nucleotidyl transferase; DUF: Domain of unknown function; ASCH: Activating signal cointegrator 1 homology.

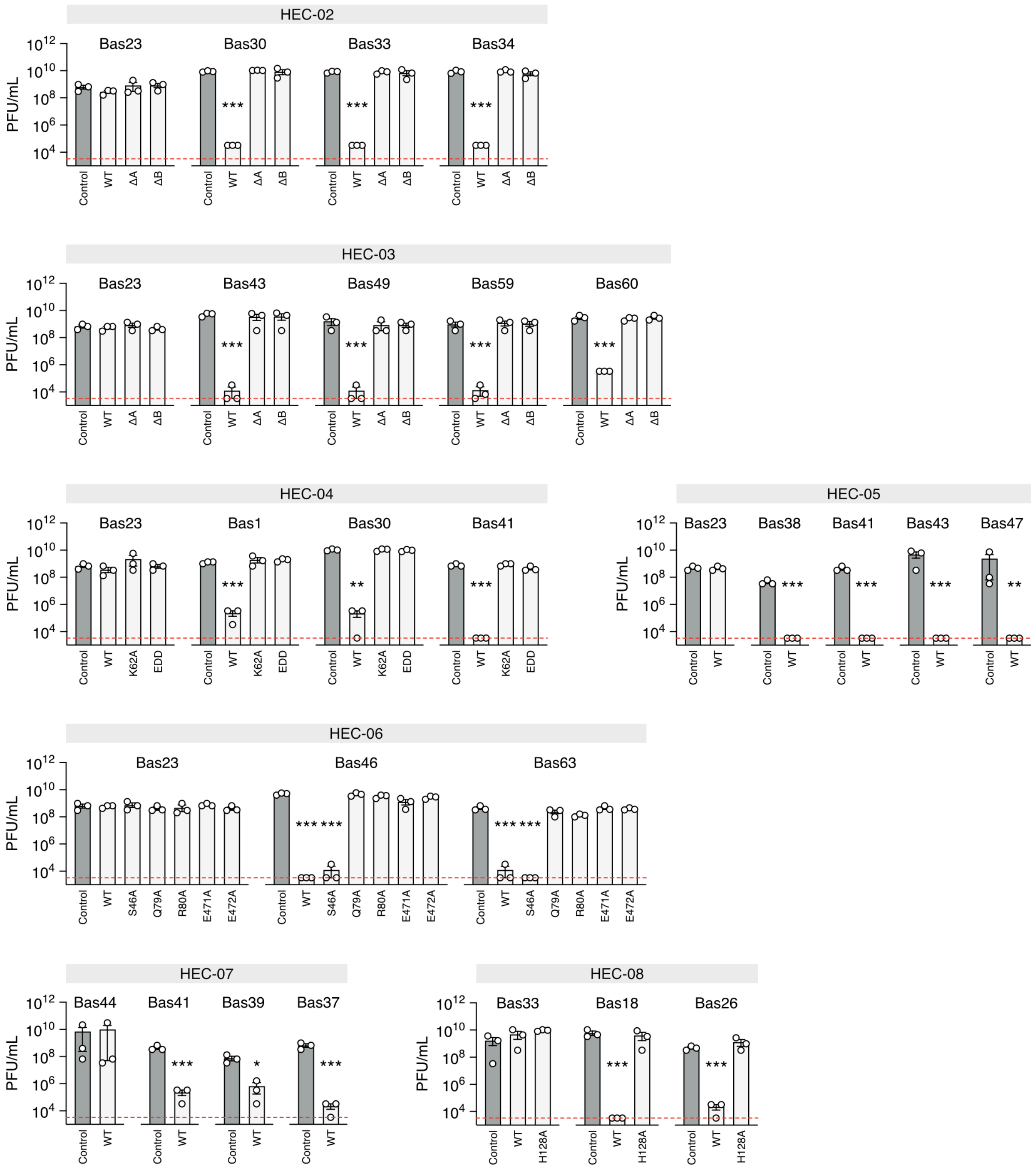


**Figure S2. Results of phage plaque assays.** Reduction of plaque forming units (PFU) data for a panel of phages with a >10-fold reduction of EOP in the presence of the respective HEC system; $n$ = 3, error bars represent the standard error of the mean. Sample means were compared by *t*-test (Student, 1908) (comparing the PFU/mL of each phage in the presence of a wildtype (WT) or mutated HEC versus the non-defence control) with Holm-Šídák correction for multiple testing (Holm, 1979); * = $p < 0.05$; ** = $p < 0.01$; *** = $p < 0.001$.

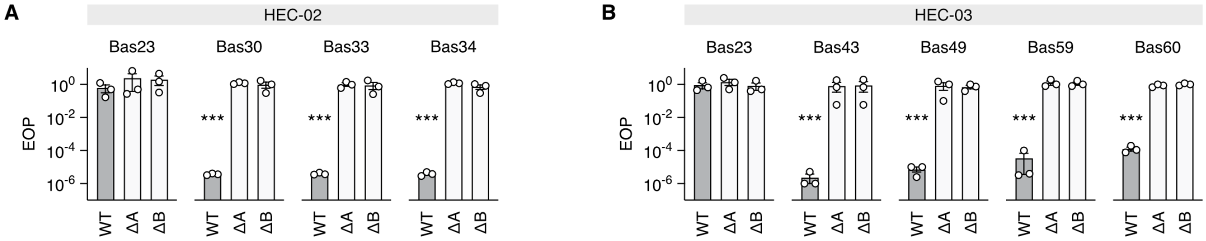


**Figure S3. HEC-02 and HEC-03 require both genes to function.** (A) Efficiency of plaquing data from mutational analysis of HEC-02. (B) Efficiency of plaquing data from mutational analysis of HEC-03. *N* = 3, error bars represent the standard error of the mean. Sample means were compared by t-test (Student, 1908) (comparing the PFU/mL of each phage in the presence of a wildtype (WT) HEC or mutated HEC versus the non-defence control) with Holm-Šídák correction for multiple testing (Holm, 1979); *** = $p < 0.001$. See **Figure S2** for assay results plotted by PFU/mL.

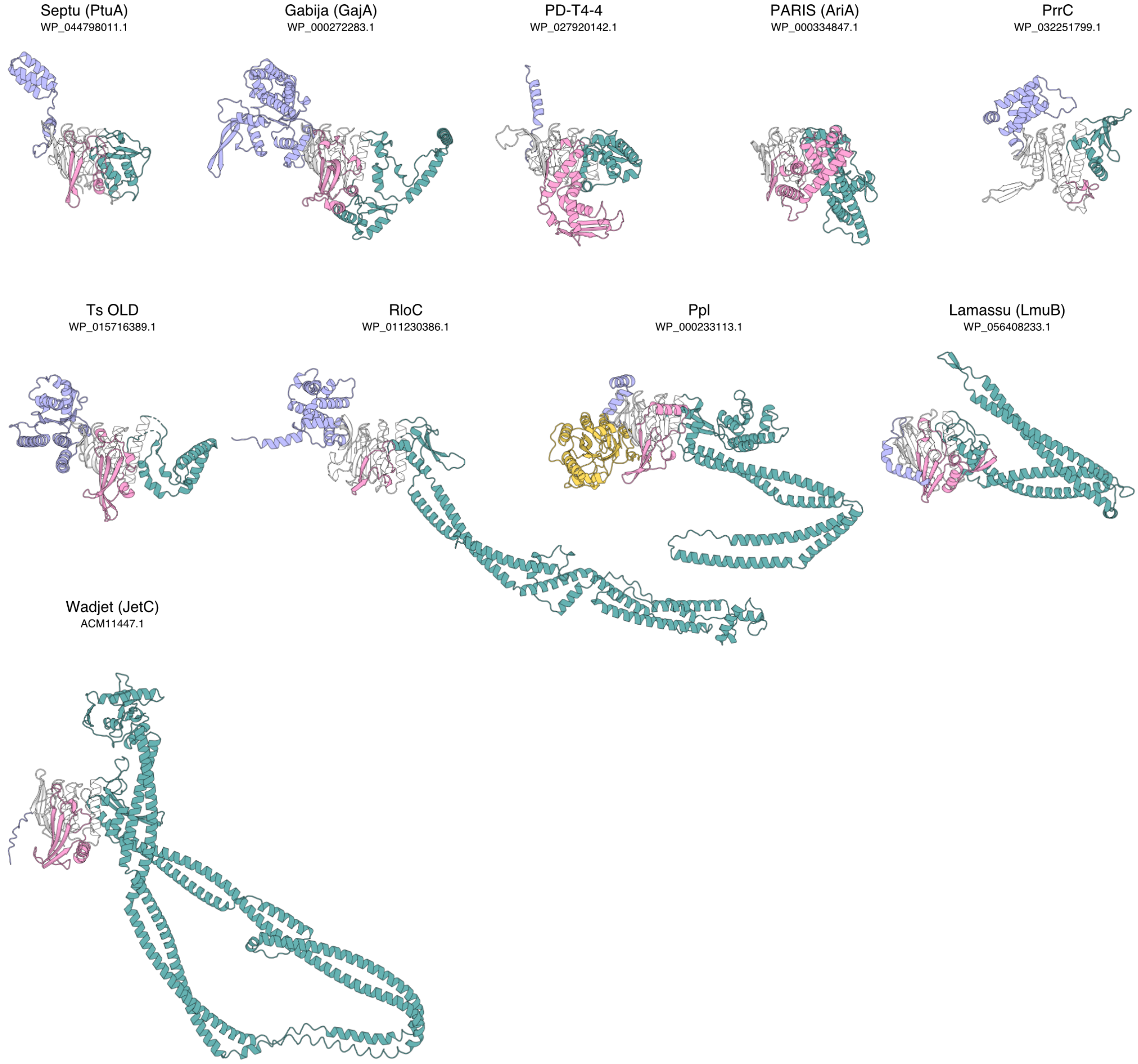


**Figure S4. ABC ATPase-based defence proteins.** Structures were predicted using ColabFold (Jumper et al., 2021; Mirdita et al., 2022). The core ATPase fold is coloured white. The first and second inserts are coloured pink and green, respectively. The N-terminal and C-terminal extensions are coloured yellow and purple, respectively.

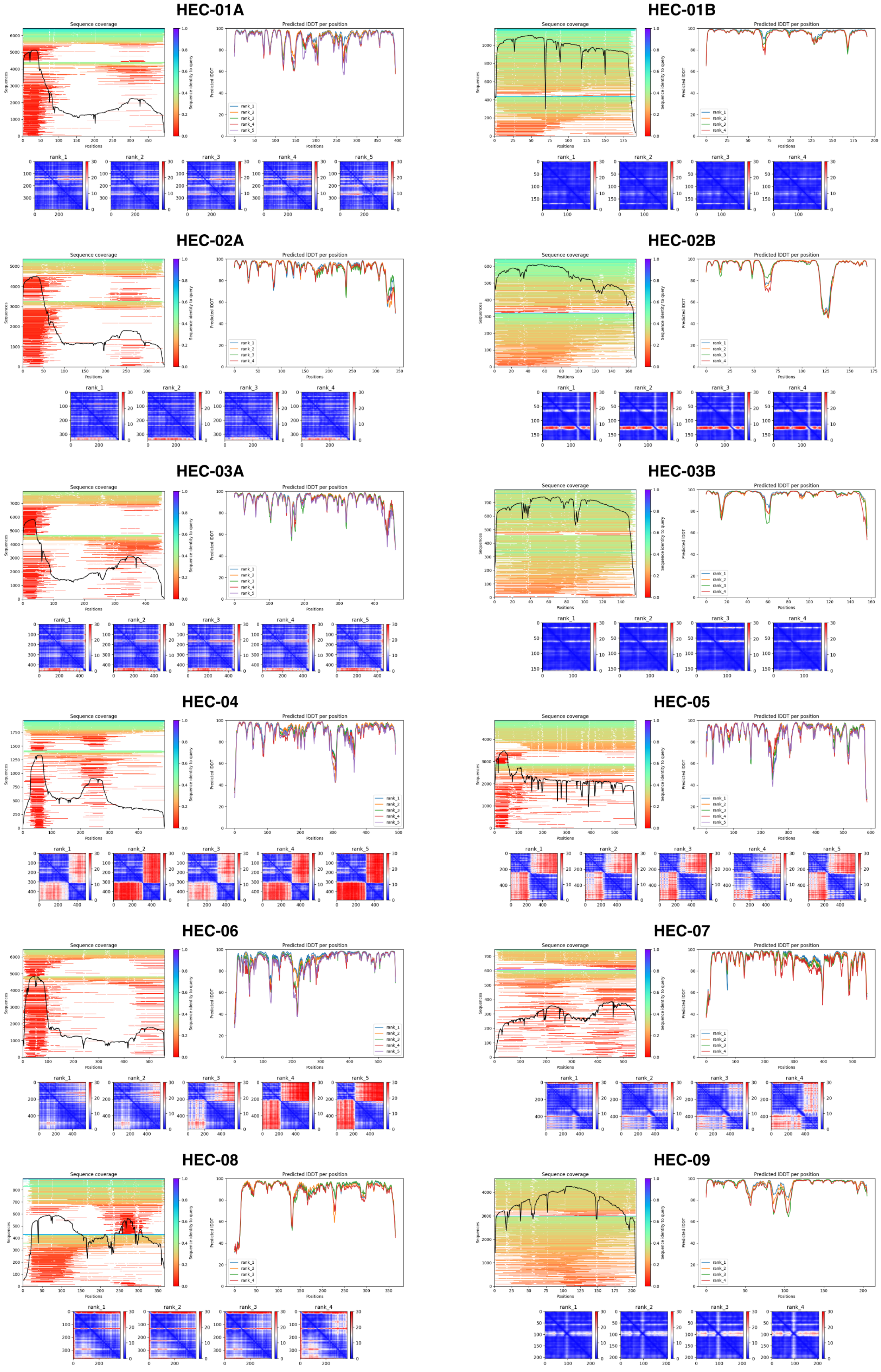


**Figure S5. Confidence metrics associated with predicted HEC structures.** The left plot shows alignment coverage during the MMseqs2 homology search step. The right plot shows the predicted local distance difference test (pLDDT), and the lower plots show the predicted aligned error (PAE).

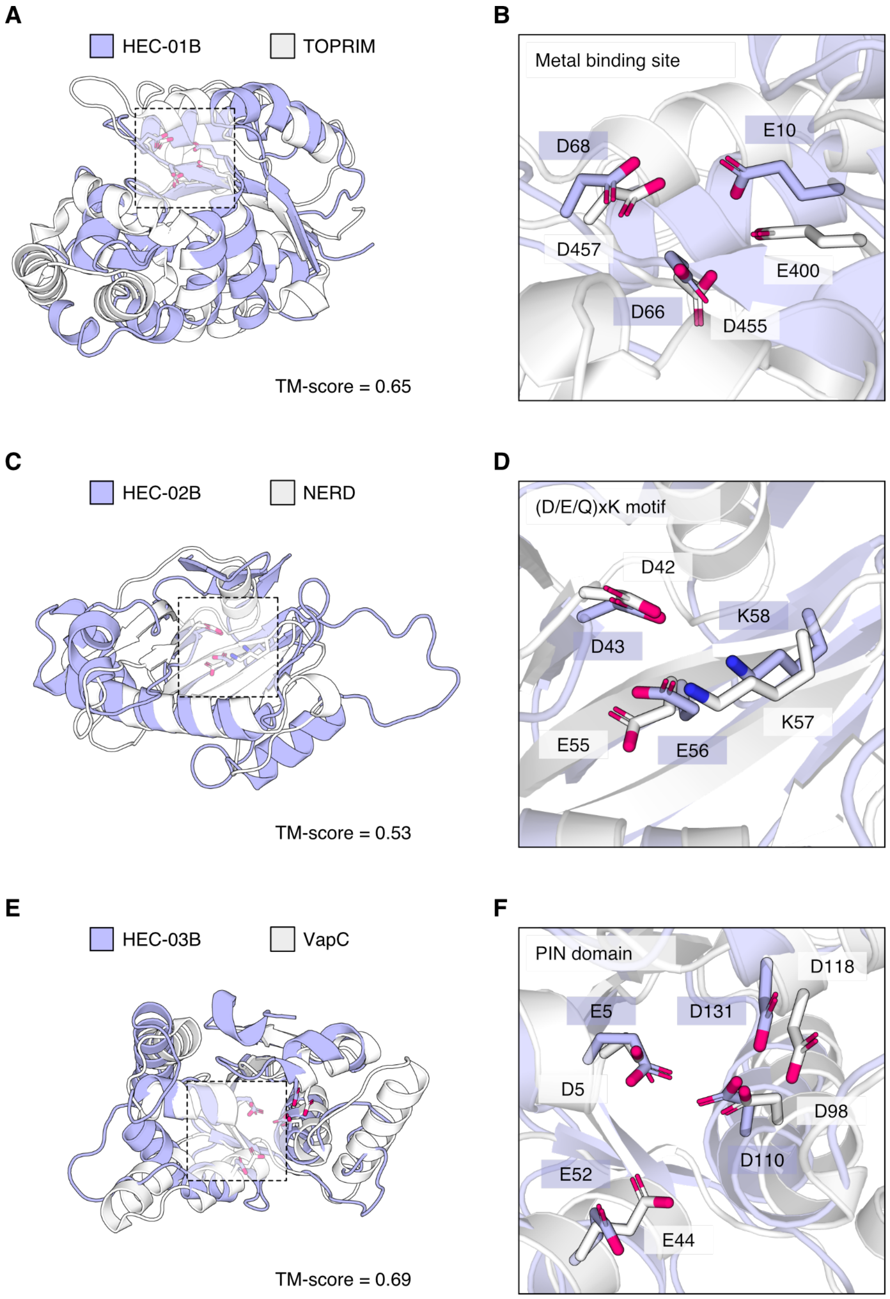


**Figure S6. HEC ABC ATPases are associated with several different putative effector proteins or domains.** (A) HEC-01B predicted structure (purple) aligned with the TOPRIM domain of Bp OLD (PDB: 6NK8; white). The conserved metal binding site presented in panel B is highlighted. (B) Conserved metal binding site residues in HEC-01B and Bp OLD. (C) HEC-02B predicted structure (purple) aligned with the Nuclease-related domain (NERD) RoseTTAFold (Baek et al., 2021) predicted structure (PFAM: PF08378; white). The conserved catalytic residues presented in panel D are highlighted. (D) Conserved catalytic residues in HEC-02B and the NERD domain. (E) HEC-03B predicted structure (purple) aligned with VapC (PDB: 6A7V; white). The shared PIN domain presented in panel G is highlighted. (F)

Conserved PIN domain in HEC-03B and VapC. TM-scores were calculated with US-align (Zhang et al., 2022), where TM-score ≥ 0.5 suggests shared global topology (Xu and Zhang, 2010).

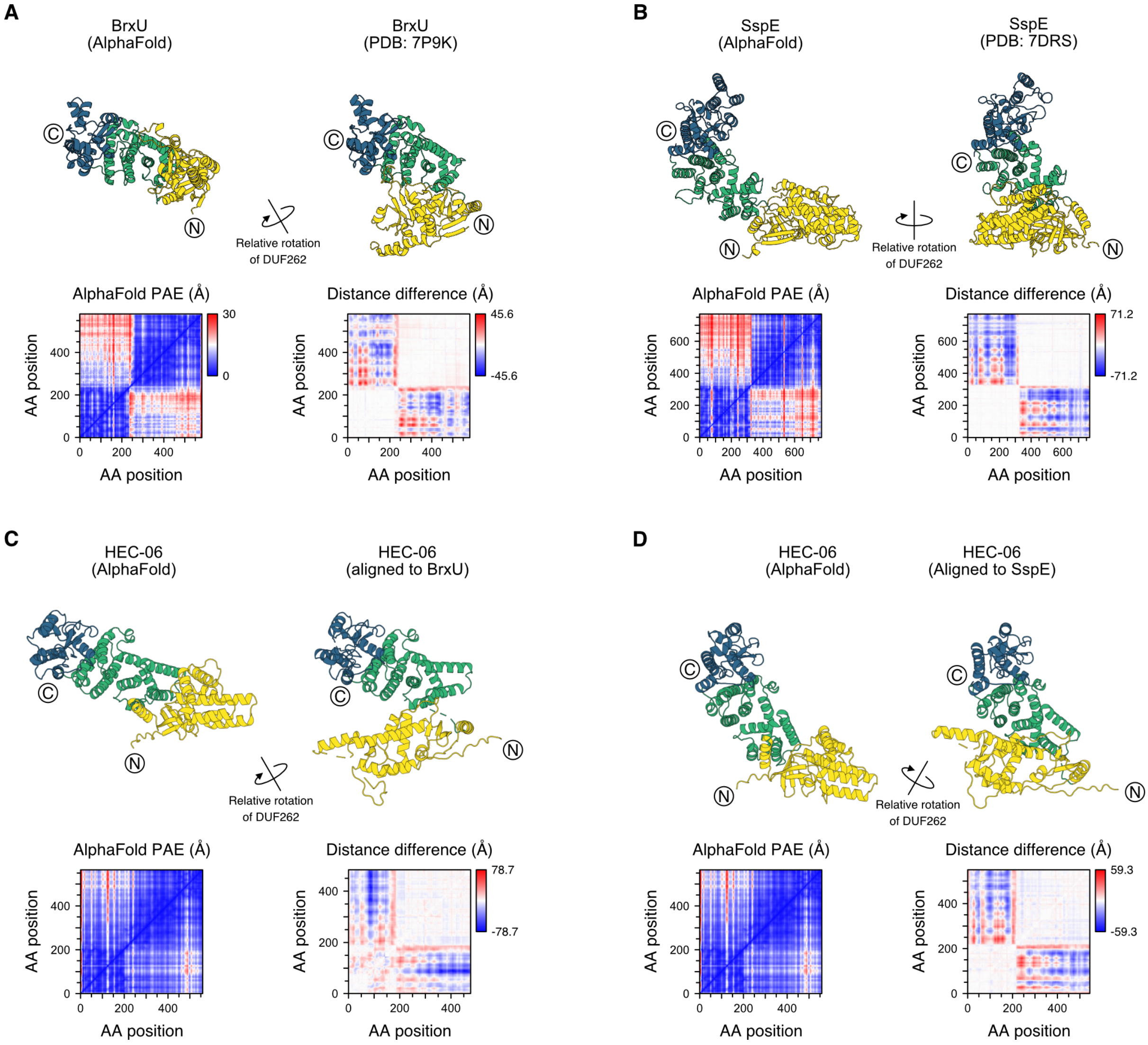


**Figure S7. AlphaFold accurately predicts the individual domains of GmrSD family proteins, but not the overall conformation.** (A) The predicted structure of BrxU (left) versus the known structure (right). Twists: 1, P-value: 0, identity: 99.5%, similarity: 99.5%. (B) The predicted structure of SspE (left) versus the known structure (right). Twists: 1, P-value: 0, identity: 97.3%, similarity: 97.5%. (C) The predicted structure of HEC-06 (left) versus a theoretical structure aligned to BrxU (right). Twists: 5, P-value: 2.12e-04, identity: 6.3%, similarity: 18.9%. (D) The predicted structure of HEC-06 (left) versus a theoretical structure aligned to SspE (right). Twists: 2, P-value: 1.01e-11, identity: 9.4%, similarity: 21.7%. The arrows show the approximate axis of rotation to twist the DUF262 domain into the expected orientation. The predicted aligned error (PAE) plots demonstrate uncertainty in the inter-domain accuracy of prediction by AlphaFold. The distance difference plots show the distance required to twist the DUF262 domain into the expected orientation, which corresponds with the uncertainty in the respective PAE plots. AA: amino acid.

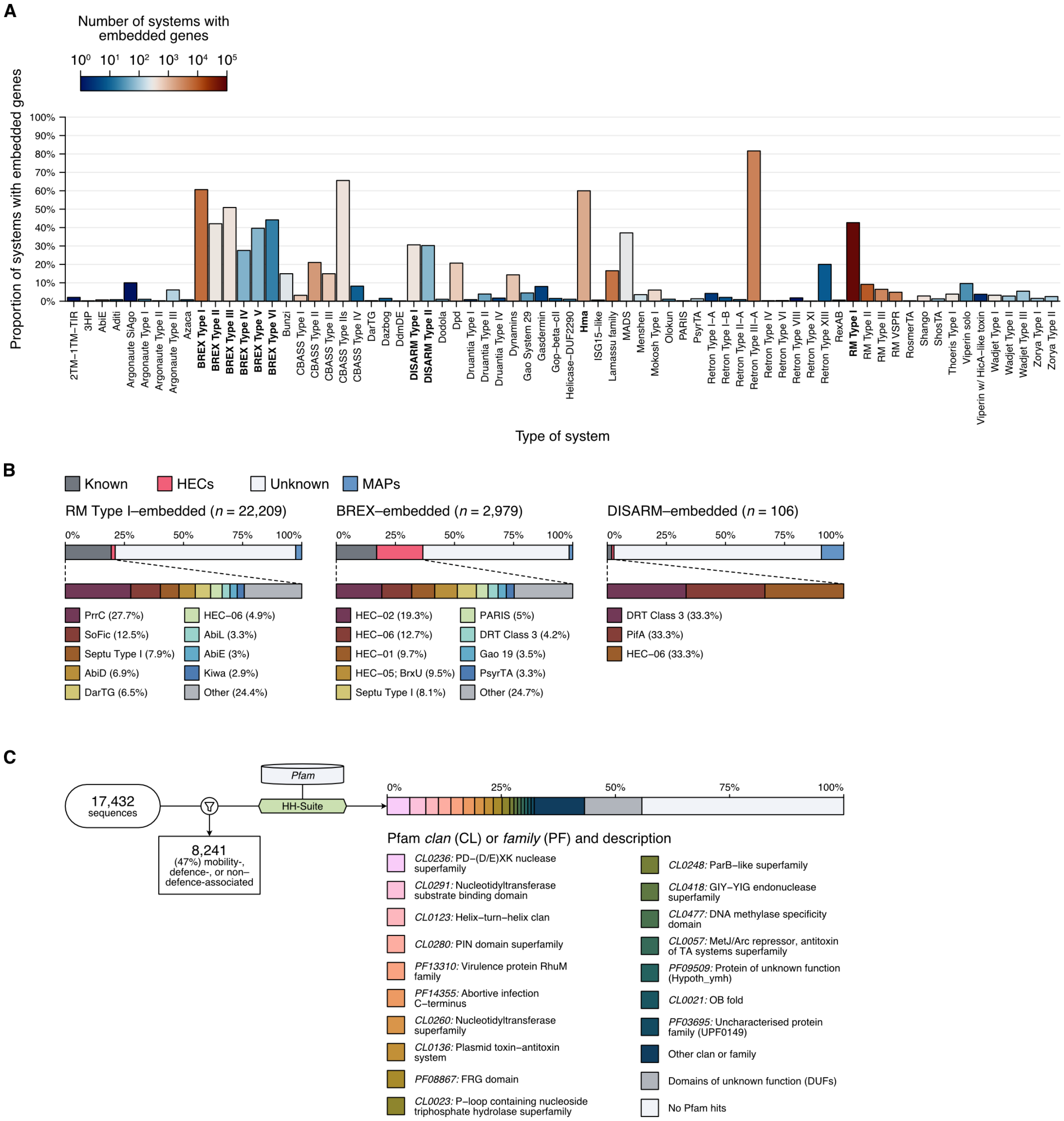


**Figure S8. Several types of defence systems often contain embedded genes.** (A) Proportion and number of defence systems that contain embedded genes. Systems with embedded genes that were investigated in this study are in bold. (B) Proportion of BREX/DISARM/RM–embedded genes identified as encoding known systems (grey, further detailed underneath), HECs (red), MAPs (blue), or otherwise unknown proteins (white). (C) Top Pfam domain annotations of BREX/DISARM/RM–embedded genes not identified as MAPs, defence-, or non–defence-associated.
