## Supplemental Table Descriptions for "New antiviral defences are genetically embedded within prokaryotic immune systems"

**Table S1:** Domain annotations for protein families embedded in more than five Hma systems.

**Table S2:** Candidate defence systems assessed for defence activity.

**Table S3:** BREX/DISARM/RM–embedded protein families versus mobileOG-db.

**Table S4:** BREX/DISARM/RM–embedded protein families versus PADLOC-DB.

**Table S5:** BREX/DISARM/RM–embedded protein families versus Defense Finder HMMs.

**Table S6:** BREX/DISARM/RM–embedded protein families versus the EggNOG database.

**Table S7:** BREX/DISARM/RM–embedded protein families versus Pfam.

**Table S8:** PDC assignments.
